# Is Evolution Predictable? Experiments in an Evolutionary Video Game

**DOI:** 10.1101/2025.11.27.691019

**Authors:** Kristen M. Martinet, Brea Dace, Luke J. Harmon, Shaelyn Pearson, John Wischnowski, Landon R. Wright, Terence Soule, Barrie D. Robison

## Abstract

The outcome of evolution sometimes appears to be predictable, as in the evolution of the same characteristics independently in convergent evolution. Other times, evolution’s path depends on starting conditions or chance events, and some forms evolve just once and never appear again. Is convergence, and by implication, predictability, a common characteristic of the evolutionary process? We aimed to answer this question by using an evolutionary video game titled *Project Hastur* as a study system. In this video game, the enemies evolve traits to help them combat the player’s strategy. We determined whether the same environmental pressures (in this case, player strategy) lead to predictable evolution. We conducted a series of experiments with three different playing styles and four different evolutionary treatments with varying kinds of selection pressure, each with several replicates. For each replicate, enemies from the first and final generations were categorized into types using k-means clustering. The Euclidean distance between the cluster centroids and the sum of squares values were recorded for each replicate and compared among evolutionary treatments using Wilcoxon rank sum tests. Evolutionary predictability was evaluated using permutation tests. We found that fitness functions in Project Hastur led to incredible, but unpredictable, diversity. The fitness landscape changes between replicates, even within the same experimental treatment and regardless of player strategy, resulting in enemies with an unpredictable array of trait values.

## Introduction

Is evolution predictable (Losos 2017)? From one point of view, the answer to this question is clearly yes; the process of evolution, involving the interactions between mutation, drift, and natural selection, can be highly repeatable, both in the lab (Rainey and Travisano 1998) and in the field (Nosil et al. 2018). Convergent evolution, where organisms evolve the same characteristics independently, has long been taken as a hallmark of the predictability of the evolutionary process (Morris 2003). If we see, for example, the same mammalian types emerging from marsupial ancestors on the continent of Australia as have arisen from placental ancestors elsewhere, then perhaps evolution is repeatable and, thus, predictable (Mazel et al. 2017). On the other hand, sometimes the outcome of evolution is highly contingent on starting conditions or chance events (Erwin 2006; Blount 2016). There are countless examples of unique organisms on Earth with forms that evolved just once. Thus, it remains unclear whether convergence – and, by extension, predictability – is a common characteristic of the evolutionary process of life on Earth (Gompert et al. 2022). Here there has been continuous and wide disagreement among scientists (Losos 2017).

Part of the disagreement stems from the mixed nature of empirical research. For example, to test for predictability, scientists have collected empirical data and compared the outcome of replicated “natural experiments” in evolution. These experiments have included a range of time spans, from the shallow (replicated evolution in stickleback over 10,000 years; Schluter 2000) to the deep (replicated radiations in Anolis lizards; Losos et al. 1998). As a whole, these studies are effective as a proof of concept: under some circumstances, predictable evolution can occur. The core problem, though, is that evidence is generally mixed, and evolution is found to be both somewhat predictable and somewhat not. For example, predictability over short time scales is limited both by the stochastic nature of evolution and the limits to the precision of our measurements of natural selection (Gompert et al. 2022). Similar problems exist over longer time scales. For example, despite extensive effort to detect them, anole ecomorphs, repeated across the four islands of the Greater Antilles (Losos et al. 1998), do not seem to exist for other island lizard groups. In fact, these ecomorphs do not even appear for anoles evolving on the mainland of South America (Patton et al. 2021). Even when considering the repeated evolution of anoles across the Greater Antilles, both phenotypic and genotypic changes are difficult to predict (Thurman et al. 2023).

Laboratory and simulation studies are also inconclusive. In both situations, it is clear that evolution can be somewhat predictable under the right conditions (Matute et al. 2020). However, under some conditions, even simulated and experimental evolution can be idiosyncratic and more difficult to predict (Milocco and Salazar-Ciudad 2020). Both lab and simulation studies suffer from a major weakness: simulation set-up and assumptions can predispose the results. For evolutionary experiments, the predictability of selection itself can determine the outcome (Lobkovsky et al. 2011). In cases where the dynamics of natural selection are predictable and repeatable, one often finds that evolutionary outcomes are the same (Rainey and Travisano 1998; Herron and Doebeli 2013). By contrast, when selection can take different paths across evolutionary replicates, then we see less predictable outcomes (Blount et al. 2008).

To overcome the limitations of both empirical and simulation studies, we need a system where one can run many replicates (as in a simulation) but where we do not have to directly specify the form of natural selection, instead allowing it to emerge, as it does in the natural world, from the interactions among organisms and their environment. For this, we turn to digital evolution. There have been several excellent implementations of digital evolution, such as Tierra (Ray 1991) and Avida (Ofria and Wilke 2004), but these popular platforms only have a “set and watch” implementation. Video games that implement digital evolution provide a more actively interactive platform for studying evolution. Creatures in video games interact with one another, their environment, and with the “player” (the person playing the video game) and these interactions determine their survival. Video game creatures exhibit variation among individuals related to survival, and many more are born than can possibly survive. One need only add inheritance of traits from parents to offspring, and these virtual creatures will evolve. Furthermore, selection in such a system emerges from the basic rules of the game, as it does in the natural world. We thus consider evolutionary video games to be a prime opportunity to evaluate the predictability of evolution and the ubiquity of predictable evolution.

In this paper, we use an evolutionary video game, called *Project Hastur* (Polymorphic Games 2019), to evaluate the predictability of evolution. Using a series of experiments with different playing styles and evolutionary treatments, we show that evolution is only predictable in terms of the relative amount of diversity produced, but that the details of the varieties that arise differ dramatically even among replicates. Among the many evolutionary outcomes we generate, one could cherry-pick particular examples of convergence - as we have, perhaps, in the natural world - but the overall picture is one of unpredictability.

## Methods

### Gameplay and Genetic Details

We developed *Project Hastur*, a tower defense style video game where enemies evolve in response to the actions of the players (Figure 1). Briefly, the player must defend their bases against waves of enemies called the “Protean Swarm” by building turrets that shoot various types of projectiles at the enemies (Proteans). Both the turrets and the enemies have distinct capabilities, strengths, and weaknesses. The unique feature of *Project Hastur* compared to other similar tower defense games is that the enemies evolve. Enemies have a genome that determines their traits, and fitness is based on their performance in attacking the player. When enemy fitness is associated with genotypes, then natural selection will lead to changes in allele frequencies within the enemy population.

**Figure 1.**
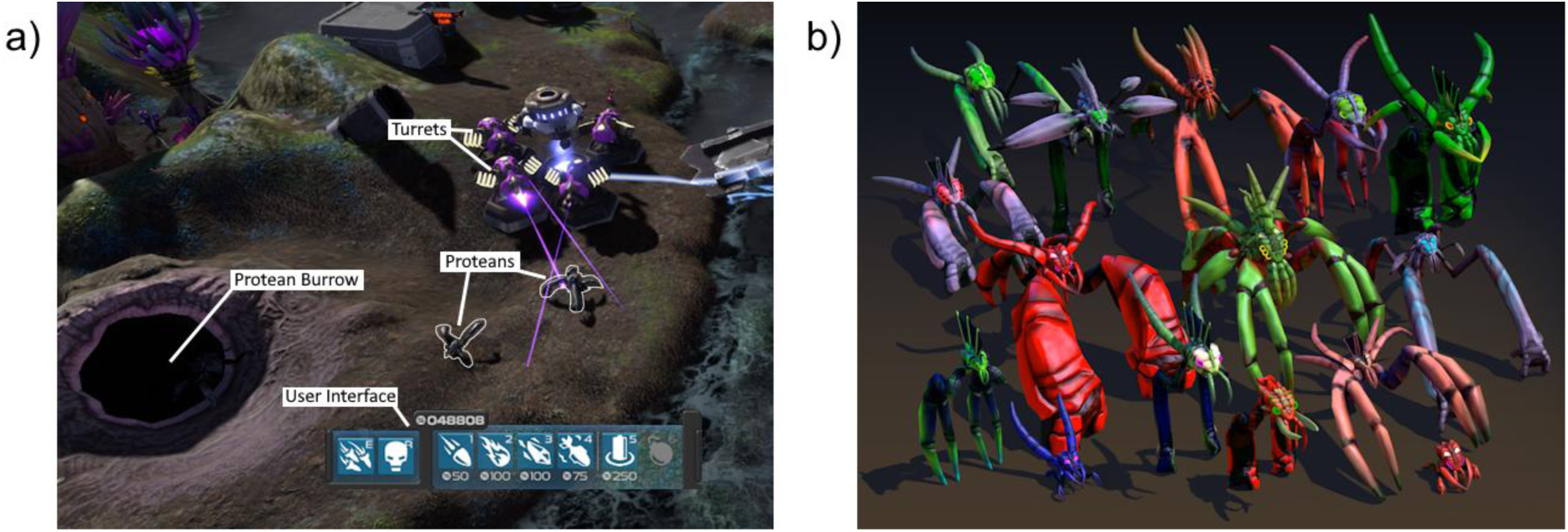
A screenshot (a) of a typical game state in Project Hastur with important features labeled, along with a sample of Protean morphological diversity (b).

Each enemy in *Project Hastur* has a set of 17 behavioral and physical traits that are determined by a diploid genome made up of 80 genes. Further details about these traits and their genetics can be found in Text S1. Some genes affect only the visual display of the enemies, but a set of genes, which we focus on here, affect Protean traits relevant to game play. Genes are not pleiotropic (each gene only affects one trait) and freely recombine. We also model mutations, with effects that follow a Gaussian distribution. In order to survive, these enemies must escape a set of turrets placed by the player. Once a turret is placed, it shoots at Proteans that walk near it. In normal (non-experimental) gameplay, the Proteans are able to destroy turrets by attacking them. Turrets come in several varieties, but the focal turret variety for this study is the kinetic type. Further details about types can be found in Text S3.

Protean enemies spawn in waves, with each wave representing a discrete generation. Proteans can reproduce either asexually or sexually. For asexual reproduction, they can clone themselves by capturing a civilian, who are humans that randomly wander around the gameplay map. Any Protean that succeeds in consuming a civilian instantly clones itself, producing a number of genetically identical offspring inversely proportional to the Protean’s size (smaller Proteans produce more). Sexual reproduction takes place between waves. After a wave, each Protean is assigned a fitness value based on its minimum distance to the human base, and, if they reached the base, how much damage they did. Proteans are then selected as parents based on their relative fitness among all candidates. As the Proteans are hermaphrodites, any two can be selected as parents, which then each pass on one allele, selected at random, for each locus in their genome. Mutations are then randomly applied to the alleles in the newly instantiated offspring’s genome.

### Data Collection

Data was collected in *Project Hastur*’s “experiment mode,” in which the player chooses turret locations that persist until the end of a designated number of generations. Further details of experiment mode parameters are included in Text S2. Different evolutionary treatments were used to evaluate the predictability of evolution and explore the impact of different fitness functions on trait space. When all fitness functions are active, Proteans can gain fitness from attacking turrets (tower fitness) and approaching human bases (base fitness). Additionally, civilians that the Proteans can use to clone themselves can be present or absent. The four evolutionary treatments are as follows: base and tower fitness inactive with civilians absent (denoted as “Control”); base and tower fitness inactive with civilians present (denoted as “OnlyClone”); base and tower fitness active with civilians absent (denoted as “OnlyFit”); and base and tower fitness active with civilians present (denoted as “FitClone”) (Figure 2). Unlike other simulations that typically specify the fitness function directly, the fitness landscape here emerges as a result of strategies employed by the player.

**Figure 2.**
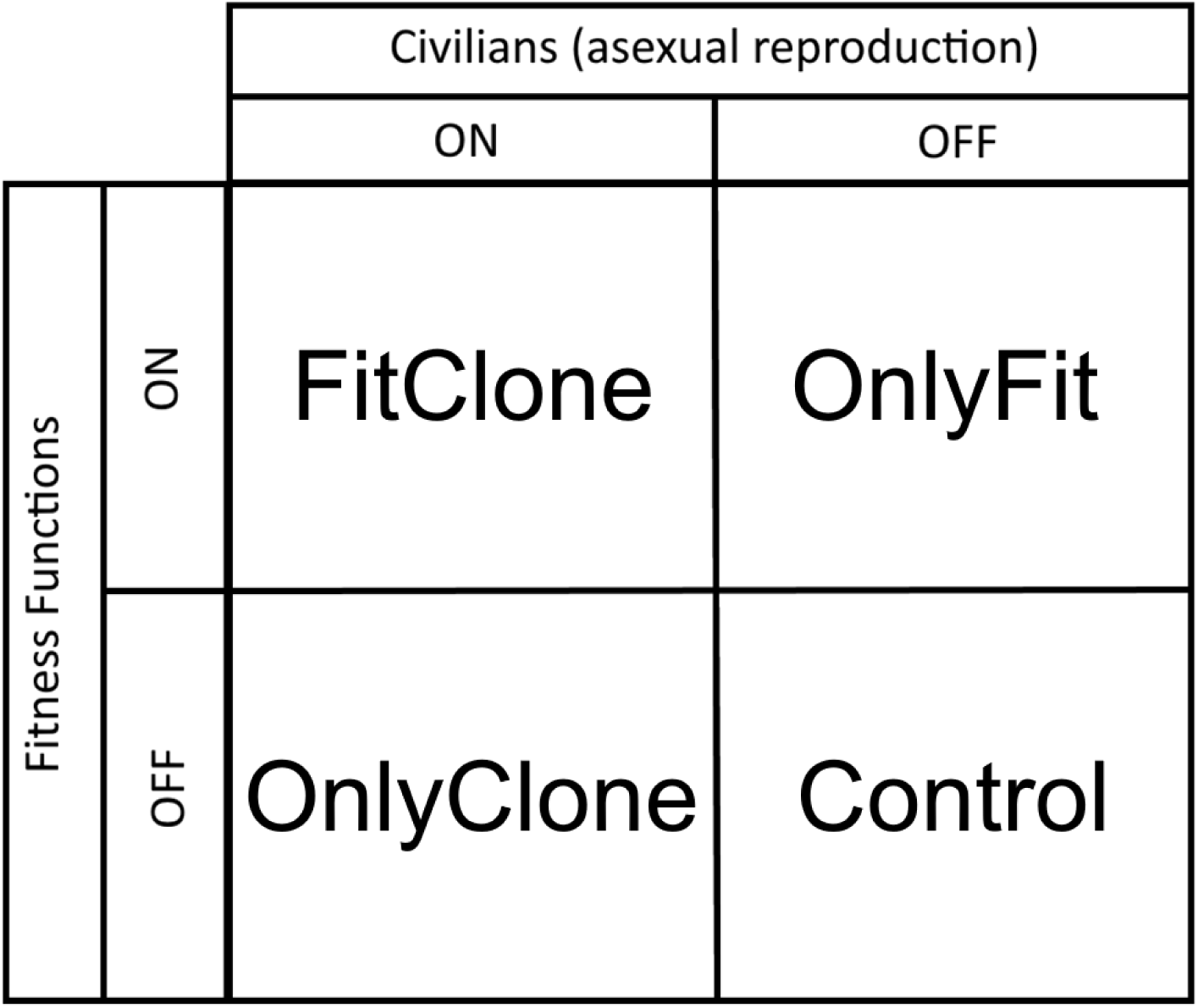
A graphical representation of the different evolutionary treatments and their fitness settings.

In addition to the evolutionary treatments, we assess the impact of player strategy on trait evolution by employing different turret arrangements. Turret arrangements were placed by a player (S.P.) using three different strategies: Burrows, Clump, and People. In the Burrows strategy, the player positioned their turrets near the burrows from which the Proteans emerge. When following the Clump strategy, the player positioned their turrets in clusters along stretches of the map that the Proteans must cross to attack the human bases. Finally, for the People strategy, the player positioned their turrets near the structures from which the civilians emerge. These turret arrangements were saved so that we could run replicates for each strategy. Each strategy was employed during each evolutionary treatment for nine replicates with 50 generations in each replicate in order to test for evolutionary predictability within treatments.

### Trait Analysis

To evaluate evolutionary predictability, we first clustered the Proteans using fourteen traits that are important for their movement and aggression (bolded traits in Table S1). These clustering analyses allow us to see how many varieties of Proteans evolved over the course of the replicate. For each replicate, trait values from the last generation were isolated. Then, model-based clustering from the R package *mclust* was used to select the best k (number of clusters) based on BIC scores (Scrucca et al. 2016). Models were estimated using expectation-maximization. Once the optimal k was selected, it was used with k-means to separate the trait space into k clusters.

These clusters served as descriptions for the different types of Proteans that evolved during each replicate. These clusters of the different types of Proteans can be compared to determine whether their evolution is predictable. In order to assess how different the Proteans from the final generation are from the first generation in each replicate, the centroids for each trait were compared to the mean trait values in the first generation using Euclidean distance. If this distance is large, then we assume that the experimental treatment promoted the evolution of Proteans that are more diverse and better-adapted to the strategy being employed compared to those found in the starting generation. If this distance is small, then we assume that the experimental treatment did not drive the evolution of diverse Proteans. In addition to comparing centroids, we also recorded the sum of squares values from the k-means clustering to test for the differences in Protean evolution within each evolutionary treatment. If the sum of squares value is large (close to 100), the variation in the replicate is captured well by the clusters and provides evidence for predictable evolution. If the sum of squares value is small, the variation in the replicate is not captured by the clusters and the evolution in that replicate is unpredictable. Finally, the distances and sum of squares values from each evolutionary treatment (FitClone, OnlyFit, OnlyClone, and Control) were compared to each other using unpaired Wilcoxon rank sum tests.

### Predictability Test

In order to assess the predictability of Protean evolution, we conducted a permutation test that assesses the similarity of evolutionary outcomes. If the replicates are predictable, we should observe that cluster centroids are unusually similar across replicates within the same evolutionary treatment. We tested this predictability by determining whether any given cluster center was closer to another center within the same experimental treatment, or if it was closer to another center from a different experimental treatment. If the Euclidean distance between centers in the same experimental treatment was smaller, we considered that result to be evidence for evolutionary predictability. For each evolutionary treatment and player strategy, one thousand of these comparisons were made against every other evolutionary treatment with the same player strategy. For each comparison, we recorded the number of times a center was closer to another center within the same experimental treatment than to a center within a different treatment. If the number of times a center was closer to another center within the same experimental treatment was greater than 95% of the permutations, we considered that to be a significant result. A significant result in this permutation test provides evidence for the presence of evolutionary predictability when the settings match the experimental treatment at hand.

## Results

The presence of fitness functions had a dramatic impact on the amount of evolutionary change we observed in each evolutionary treatment, and player strategy did not have a specific impact in any evolutionary treatment (Table 1). When comparing the distances for each evolutionary treatment to each other using a Wilcoxon rank-sum test, we expected that treatment FitClone would have significantly greater distances than the Control, regardless of strategy, because FitClone includes all fitness functions and the Control includes no fitness functions. Additionally, we expected that the Control would never have significantly greater distances compared to the other evolutionary treatments because the other three treatments have at least one active fitness function. These two expectations were met regardless of strategy (Table 1).

**Table 1.** Wilcoxon rank-sum p-values for comparing Euclidean distances between the mean trait values from the first generation and the centroids from the last generation within each evolutionary treatment to the distances within every other evolutionary treatment. The first two columns of each row signify which evolutionary treatments are being compared in the Wilcoxon rank-sum test with the equation Focal Treatment > Comparison. Bolded p-values are significant (α = 0.05).

| Focal Treatment | Comparison | Player Strategy |  |  |
| --- | --- | --- | --- | --- |
|  |  | Burrow | Clumped | People |
| FitClone | Control | **2.06E-05** | **2.06E-05** | **2.06E-05** |
| FitClone | OnlyClone | **2.06E-05** | **2.06E-05** | **2.06E-05** |
| FitClone | OnlyFit | 0.3652 | 0.2181 | 0.1290 |
| Control | FitClone | 1 | 1 | 1 |
| Control | OnlyClone | 0.8513 | 1 | 0.3652 |
| Control | OnlyFit | 1 | 1 | 1 |
| OnlyClone | FitClone | 1 | 1 | 1 |
| OnlyClone | Control | 0.1701 | **4.11E-05** | 0.6668 |
| OnlyClone | OnlyFit | 1 | 1 | 1 |
| OnlyFit | FitClone | 0.6668 | 0.8067 | 0.8888 |
| OnlyFit | Control | **2.06E-05** | **2.06E-05** | **2.06E-05** |
| OnlyFit | OnlyClone | **2.06E-05** | **2.06E-05** | **2.06E-05** |

We found that the impact of asexual reproduction on fitness resulted in a considerably smaller effect on diversity compared to base and tower fitness. The presence of asexual reproduction during an experimental treatment affects the composition of the final population of Proteans, but we expected that this ability to clone would have a much smaller impact on Protean diversity compared to the presence of fitness functions. This expectation is met, as demonstrated by the comparisons between OnlyClone and the other experimental treatments. OnlyClone has significantly greater distances than the Control once, when the player utilizes the Clumped strategy, but it otherwise does not have significantly greater distances than any other experimental treatment (Table 1). To further demonstrate the smaller impact that cloning has on the Protean population compared to the fitness functions, OnlyFit and FitClone never have a significantly greater distance compared to the other, regardless of the directionality of the comparison (Table 1). This lack of significance meets our expectations because fitness is the main driver of Protean diversity in these experimental treatments, and the presence of asexual reproduction has a minimal impact.

To further support the finding that the presence of fitness functions leads to more diverse Proteans by the final generation, we compared sum of squares values from the k-means clustering between the evolutionary treatments. Similarly to comparing the distances between the mean trait values in the first generation and the centroids from the final generation, we expected that evolutionary treatment FitClone would have higher sum of squares values compared to all the other treatments. We also expected that the Control would never have larger sum of squares values compared to the other treatments because the lack of fitness functions leads to a much less diverse Protean population. Both of these expectations are met (Table 2).

**Table 2.** Wilcoxon rank-sum p-values for comparing sum-of-squares values within each evolutionary treatment to every other evolutionary treatment. The first two columns of each row signify which evolutionary treatments are being compared in the Wilcoxon rank-sum test with the equation Focal Treatment > Comparison. Bolded p-values are significant (α = 0.05).

|  |  | Player Strategy |  |  |
| --- | --- | --- | --- | --- |
| Focal Treatment | Comparison | Burrow | Clumped | People |
| FitClone | Control | **0.007096** | **4.11E-05** | **2.06E-05** |
| FitClone | OnlyClone | **0.003887** | **0.000617** | **4.11E-05** |
| FitClone | OnlyFit | 0.3981 | 0.3652 | 0.2181 |
| Control | FitClone | 0.9947 | 1 | 1 |
| Control | OnlyClone | 0.2729 | 0.9433 | 0.2181 |
| Control | OnlyFit | 1 | 1 | 0.8067 |
| OnlyClone | FitClone | 0.9972 | 0.9996 | 1 |
| OnlyClone | Control | 0.7553 | 0.06796 | 0.2181 |
| OnlyClone | OnlyFit | 1 | 0.9999 | 0.9999 |
| OnlyFit | FitClone | 0.6348 | 0.6668 | 0.8067 |
| OnlyFit | Control | **2.06E-05** | **2.06E-05** | **8.23E-05** |
| OnlyFit | OnlyClone | **2.06E-05** | **0.000247** | **0.000144** |

As seen in the distance comparisons, we expected that the ability to clone would not result in a more diverse Protean population overall. This expectation is met, as evolutionary treatment OnlyClone never has larger sum of squares values than any other experimental treatment (Table 2). Finally, when comparing evolutionary treatment OnlyFit to the other treatments, we expected that OnlyFit’s sum of squares values would be greater compared to those from the Control and OnlyClone, but not when compared to those from FitClone. This expectation is met, which further solidifies the fact that fitness functions are the most instrumental factor in the creation of diverse Protean populations. Evolutionary treatments FitClone and OnlyFit result in very similar sum of squares values, and both result in much greater sum of squares values compared to evolutionary treatment OnlyClone and the Control (Figure 3). The presence of Protean diversity is strongly dependent upon the presence of fitness functions and is not dependent on the ability for the Proteans to reproduce asexually or changes in player strategy.

**Figure 3.**
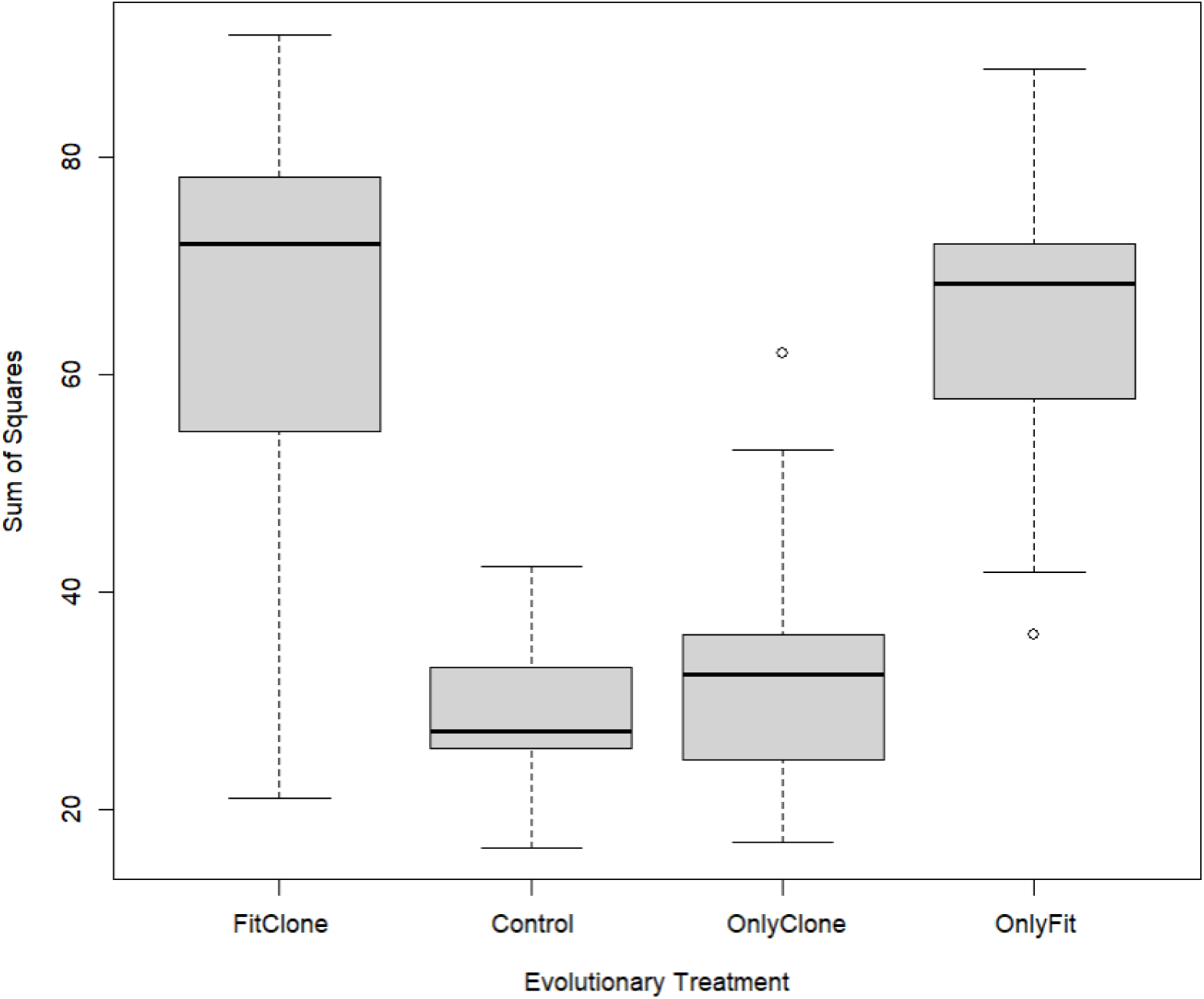
Box plot comparing the sum of squares values for each evolutionary treatment. Sum of squares values differ across evolutionary treatments (ANOVA p-value = 2.02E-23). Treatments with fitness functions active (FitClone and OnlyFit) have substantially larger sum of squares values compared to evolutionary treatments with fitness functions inactive (Control and OnlyClone).

The Wilcoxon rank-sum tests provide evidence for increased Protean diversity by the final generation, but they do not address our question of evolutionary predictability. To determine whether evolution is predictable in any experimental treatment, we conducted permutation tests amongst the experimental treatments. When using permutation tests to assess the predictability of Protean evolution, we found that none of the experimental treatments had a significant number of cluster centers that were closer to each other compared to the Euclidean distance between the given center and a center from a different experimental treatment (Table 3). While active fitness functions certainly result in a greater diversity of Proteans, the permutation tests reveal that the exact nature of this diversity is unpredictable regardless of the strategy the player uses. When fitness functions are inactive, as in the case of the Control and OnlyClone treatments, random drift results in unpredictable evolution. When fitness functions are active, as in the FitClone and OnlyFit treatments, the fitness landscape varies dramatically between replicates because of the unique emergence of different fitness peaks to combat player strategy. These differences between replicates result in unpredictable evolution when fitness functions are active as well.

**Table 3.** The proportion of permutations in which Euclidean distances between cluster centers in the focal treatment are closer to each other than to cluster centers in the comparison treatment. None of the proportions are significant.

| Focal Treatment | Comparison | Player Strategy |  |  |
| --- | --- | --- | --- | --- |
|  |  | Burrow | Clumped | People |
| FitClone | Control | 0.695 | 0.760 | 0.765 |
| FitClone | OnlyClone | 0.713 | 0.791 | 0.801 |
| FitClone | OnlyFit | 0.519 | 0.503 | 0.360 |
| Control | FitClone | 0.626 | 0.767 | 0.710 |
| Control | OnlyClone | 0.601 | 0.938 | 0.358 |
| Control | OnlyFit | 0.841 | 0.908 | 0.590 |
| OnlyClone | FitClone | 0.504 | 0.533 | 0.817 |
| OnlyClone | Control | 0.459 | 0.686 | 0.582 |
| OnlyClone | OnlyFit | 0.763 | 0.731 | 0.753 |
| OnlyFit | FitClone | 0.400 | 0.402 | 0.609 |
| OnlyFit | Control | 0.613 | 0.529 | 0.664 |
| OnlyFit | OnlyClone | 0.749 | 0.581 | 0.661 |

## Discussion

As an evolutionary video game, *Project Hastur* has unique properties that allow us to evaluate the predictability of evolution in a way that is impossible with traditional evolutionary simulations. For example, simulation studies often must determine a selection function beforehand, choose a specific fitness landscape, and select a particular set of starting generation parameters (e.g. choosing a starting population from the Freezer in Avida; Pennock 2019), but the user does not make these choices in Project Hastur. While the relative amount of variation within the Protean population was predictable, the morphologies of successful Proteans were not predictable in any of the evolutionary treatments. When given the right conditions, Proteans evolve a wide range of morphological traits that allow them to combat the player’s strategies (Figure 4). The evolution of diversity itself is predictable, although only when certain criteria are met. Within these diverse populations, however, we cannot predict how Proteans will differ from each other.

**Figure 4.**
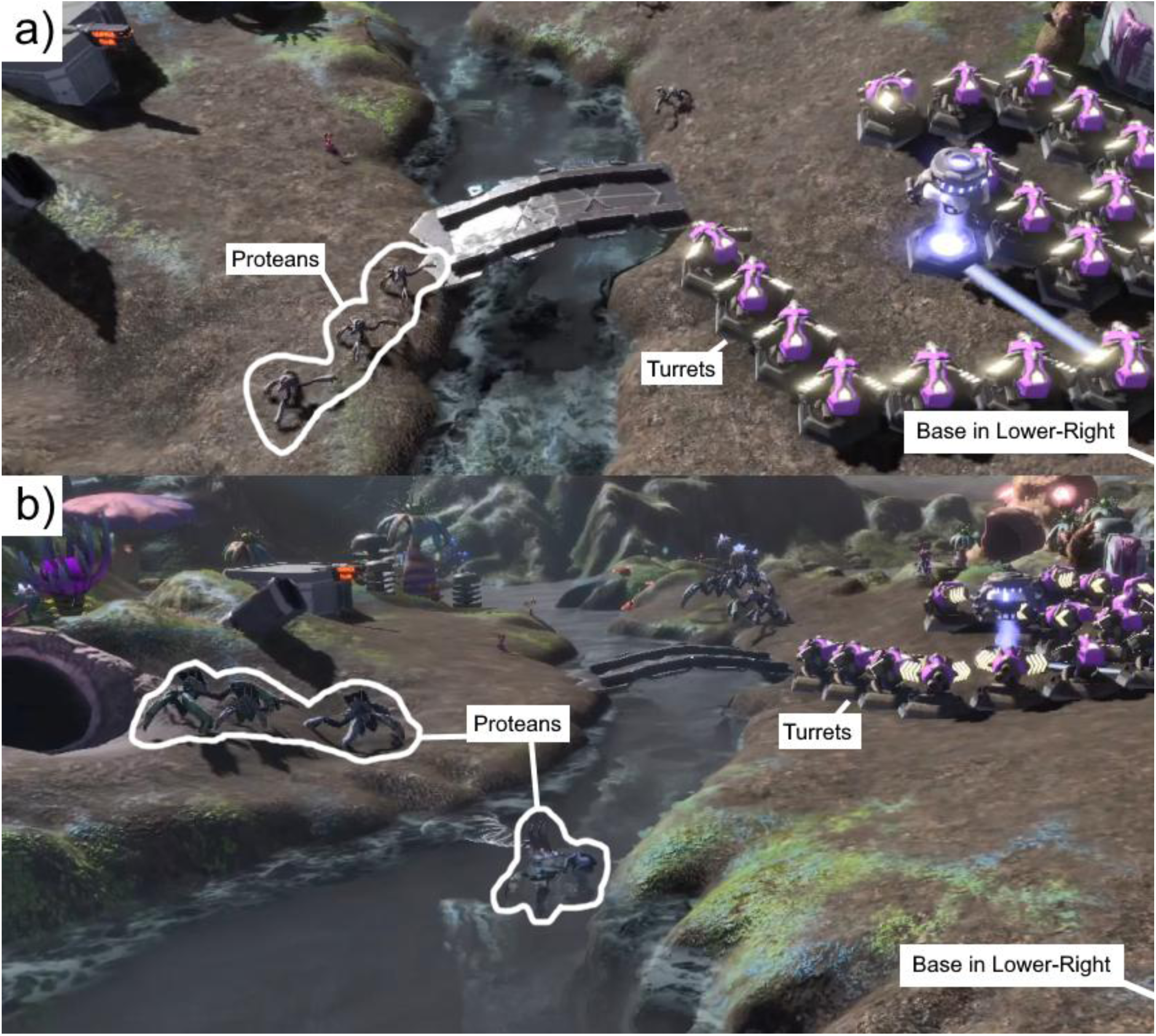
An example of Protean evolution to combat player strategy. In the first generation (a), Proteans are unable to cross the river and must walk across the bridge and through a gauntlet of turrets to get to the base. By the fourteenth generation (b), some Proteans have evolved enhanced swimming abilities that allow them to cross the river and avoid the turret gauntlet to reach the base.

Protean diversification is driven almost entirely by the presence of fitness functions, and the ability for Proteans to asexually clone themselves has a small effect on the resulting diversity of the final generation. In experimental treatments with fitness functions active (FitClone and OnlyFit), the resulting Protean populations are much more diverse than in experimental treatments with fitness functions inactive (OnlyClone and Control). During experimental treatment OnlyClone, the only driver of diversity is the ability for Proteans to asexually reproduce. OnlyClone never has significantly greater distances than any other experimental treatment, except for one comparison with the Control (Table 1). If asexual reproduction contributed greatly to the overall diversity of the Protean population, OnlyClone would have significantly greater distances than Control in all cases. In addition, OnlyFit and FitClone never have a significantly greater distance compared to the other regardless of directionality (Table 1). These comparisons also demonstrate that asexual reproduction does not contribute enough to Protean diversity to create significantly larger distances in experimental treatment FitClone.

One reason for the lack of predictability in Project Hastur might come from the unpredictable nature of selection itself. The fitness landscape changes between replicates, even within the same experimental treatment, which results in populations of Proteans that have an unpredictable array of trait values. Player strategy certainly contributes to the diversity of Proteans, but evolution responds to that strategy differently in every replicate. When we attempted to create dendrograms that visually represent the types of Proteans that emerge during each replicate, naming the tips proved to be a nonsensical task. Fourteen traits contribute to the clustering of Protean types, and attempting to incorporate all of them into names for each type of Protean resulted in uninformative names that were rarely applicable to more than one replicate (see Text S4 for more information). The changing nature of selection in Project Hastur might have impacted our ability to predict evolution, even though replicates should reduce uncertainty (Gompert et al. 2022; Nosil et al. 2020). Additionally, the high dimensionality of Protean trait space could have introduced chaotic dynamics into their evolution that prevented predictability (Doebeli and Ispolatov 2014).

The unique aspect of Project Hastur as an evolutionary video game gives our inability to predict Protean evolution particular relevance. In more traditional simulations, the fitness landscape is often determined at the beginning of the simulation through parameter choice. In Project Hastur, the fitness landscape changes between replicates even if the experimental settings remain the same. Our initial assumption while developing Project Hastur was that the fitness landscape would change between game sessions because the player and the Proteans would be in an evolutionary arms race, with Proteans combatting the player’s strategies through evolution and the player responding by changing their strategies. However, the turret arrangement for the entirety of each 50-generation replicate corresponding to a particular strategy was exactly the same, and strategy ultimately did not drive predictable evolution of Proteans (Table 3). More analysis of the fitness landscapes that emerge during a Project Hastur replicate is needed to disentangle the potential role of player strategy in the evolution of Proteans.

The unpredictable nature of evolution in Project Hastur may also be a result of the interaction between each Protean’s random starting position on the fitness landscape and the rugged shape of the fitness landscape. Once a Protean evolves to find an adaptive peak, it often cannot evolve to move beyond that peak. The different clusters that emerge from each replicate are representative of multiple fitness peaks that the Protean population found, and since the fitness landscape changes with every replicate, the accessible fitness peaks also change. This combination can result in epistasis, where groups of Proteans within a population follow evolutionary paths that cannot be reversed. Project Hastur provides a unique opportunity to study epistasis as an emergent property of the video game. These complex interactions, in turn, may be the main explanation for the fundamental failure to observe replicated radiations in our study.

The results presented here only scratch the surface of the possible evolutionary explorations that could be conducted with Project Hastur. We used one player’s data for the analyses here, but comparing data across different players would provide us with the opportunity to ask more questions about predictability. Are replicates more predictable when conducted by a player with a different interpretation of any given strategy? Does the same amount of diversity emerge from different implementations of each strategy? There are also two additional kinds of turrets available in Project Hastur (Flamethrower and Chip Shredder) that could impact the evolution of Proteans in unique ways not explored here. Do Proteans evolve different or more predictable strategies when encountering different types of turrets? Project Hastur’s Experiment Mode allows the player to turn on subpopulations, where each Protean burrow represents an independently evolving subpopulation of enemies. If subpopulations are active, is evolution more predictable? Would replicated radiations emerge from the subpopulations?

Returning to our opening question: is evolution predictable (Losos 2017)? We cannot provide a definitive answer for every instance of evolution using a video game. Still, our results - where fitness emerges naturally as a consequence of the struggle to survive and reproduce in a complex system - give some insights into evolutionary predictability. In this case, we find that only diversity itself is predictable. We can identify the conditions under which we will produce a diversity of forms. The individual identity of those forms, however, varies dramatically from one replicate to the next. Our simulations are reminiscent in some ways of broad comparisons that can be made across the Earth. If one was to search our simulation results, one might occasionally find strikingly similar forms from independent replicates - just as one can sometimes find spectacular examples of convergent evolution across continents. However, on the whole, our evolutionary outcomes were highly variable and unpredictable even across replicates with the same conditions. We suggest that the lack of predictability in our study is more likely a result of the complex interactions among genes, traits, and fitness that emerge during gameplay, and that these interactions create a system that is predictable only in the extent, but not the types, of diversity that forms.

## Code Availability

*Project Hastur*, which includes the Experiment Mode used for data collection here, is freely available on Steam: https://store.steampowered.com/app/800700/Project_Hastur/.

## Acknowledgements

We thank members of the Project Hastur development team who did not explicitly contribute to this research, but without whom this work would not exist: Samantha Heck, Patrick Vanvorce, Nicholas Valenti, Adam Odell, Cameron Perry, Tristan Lassiter, Courtney Bryant, Emily Ball, Samuel Carlson, Weston Durland, Marlan Smith, and Jacob Robison. KM was supported by the University of Idaho Institute for Interdisciplinary Data Sciences (IIDS), Bioinformatics and Computational Biology Program (BCB), and Department of Biological Sciences.

