## Supplemental Text for "Is Evolution Predictable? Experiments in an Evolutionary Video Game"

### In this document

**Text S1**. Additional Genetic Details

**Text S2**. Experiment Mode Details

**Text S3**. Additional Game Information

**Text S4**. Dendrogram Creation Details

**Text S1. Additional Genetic Details**

Proteans have 17 traits that control their behavior and morphology, described in Table S1. Traits in bold are those that we analyzed in the main text of this paper. Most trait values are calculated using sigmoid functions with bounds determined by variables that are instantiated when the game mode is selected (known as "CreatureParameters"). These values are constant throughout Experiment Mode replicates and are listed in Table S2. Trait values for Attraction0, Attraction1, and Attraction2 are simply their raw genetic values with no sigmoid or matrix transformation. The genetic values for these attraction-related genes start at 10 for Attraction0, 9 for Attraction1, and 11 for Attraction2. These starting values are the same for every Protean and were chosen to balance the gameplay. These attraction-related genes still evolve in the same way as the rest of the genome. The trait value for Collider Surface Area is calculated by taking the sum of the surface areas of all of the colliders present on the Protean. Colliders are used in the Unity Engine to determine the shape and size of an object so that the game knows when certain objects (for example, bullets from turrets) are touching other objects (for example, a Protean's body). Collider Surface Area serves as an accurate measurement of the overall size of the Protean.

| Trait | Definition |
| --- | --- |
| **ACCELERATION** | The rate at which the Protean can transition from WALK SPEED to RUN SPEED. |
| ACID RESIST | Determines the proportion of acid damage that does not affect HEALTH. |
| **ARMOR** | Determines the probability of a physical attack affecting HEALTH. |
| **ATTRACTION0** | Indicates the strength of behavioral preference (negative values) or aversion (positive values) for the player's towers. |
| **ATTRACTION1** | Indicates the strength of behavioral preference (negative values) or aversion (positive values) for civilians on the map. |
| **ATTRACTION2** | Indicates the strength of behavioral preference (negative values) or aversion (positive values) for the player's base. |
| **COLLIDER SURFACE AREA** | The total surface area of all physics colliders (defined by game engine) for the Protean. The best proxy for body size. |
| **DAMAGE** | The amount of health removed from a player structure or civilian when attacked by the Protean. |
| FIRE RESIST | Determines the proportion of fire damage that does not affect HEALTH. |
| **HEALTH** | The total amount of damage a Protean can receive before it is killed. Often referred to as "Hit Points" in video games. |
| ICE RESIST | Determines the proportion of ice damage that does not affect HEALTH. |
| **JUMP** | A threshold trait that allows Proteans to exploit jump points in each map. |
| **RUN SPEED** | The maximum travel speed of the Protean. Proteans transition from WALK SPEED to RUN SPEED at the rate defined by ACCELERATION. This transition is triggered by damage or identification of ATTRACTION game objects within its SIGHT RANGE. |
| **SIGHT RANGE** | The sensory radius of the Protean. Detects all objects of the type tower, civilian, and base within the radius. |
| **SWIM** | A threshold trait that allows Proteans to exploit swim points in each map. |
| **TURN RATE** | The rate at which the Protean can rotate its orientation. The best game proxy for agility. |

| | |
|---|---|
| **WALK SPEED** | The base movement speed for the Protean. Proteans begin each wave at WALK SPEED, and can only transition to RUN SPEED under specific conditions. |

Table S1. Definitions for the traits used in the Protean genome. Traits used in the analyses in the main text are bolded.

| Trait | x | y | z |
|---|---|---|---|
| Acceleration | 1 | 15 | 45 |
| Acid Resist | 0 | 0 | 1 |
| Armor | 0 | 0.05 | 0.5 |
| Damage | 25 | 50 | 400 |
| Fire Resist | 0 | 0 | 1 |
| Health | 250 | 1000 | 60000 |
| Ice Resist | 0 | 0 | 1 |
| Jump | 0 | 1 | 3 |
| Run Speed | 10 | 20 | 45 |
| Sight Range | 5 | 10 | 30 |
| Swim | 0 | 1 | 3 |
| Turn Rate | 240 | 360 | 720 |
| Walk Speed | 5 | 7 | 15 |

Table 2. The “CreatureParameters” that determine the bounds of the functions that are used to calculate Protean trait values. The “x” column represents the lower bound, the “y” column represents the center, and the “z” column represents the upper bound. Attraction0, Attraction1, Attraction2, and Collider Surface Area are not included in this table because they are calculated differently (see Text S1).

**Text S2. Experiment Mode Details**

*Project Hastur* includes an Experiment Mode, which allows the user to modify game states, parameters of the evolutionary model, and conditions related to running replicates (Figure S1). We used Experiment Mode to run all of the replicates for the main text. A detailed list of game parameters that can be adjusted in Experiment Mode can be found in Table S3.

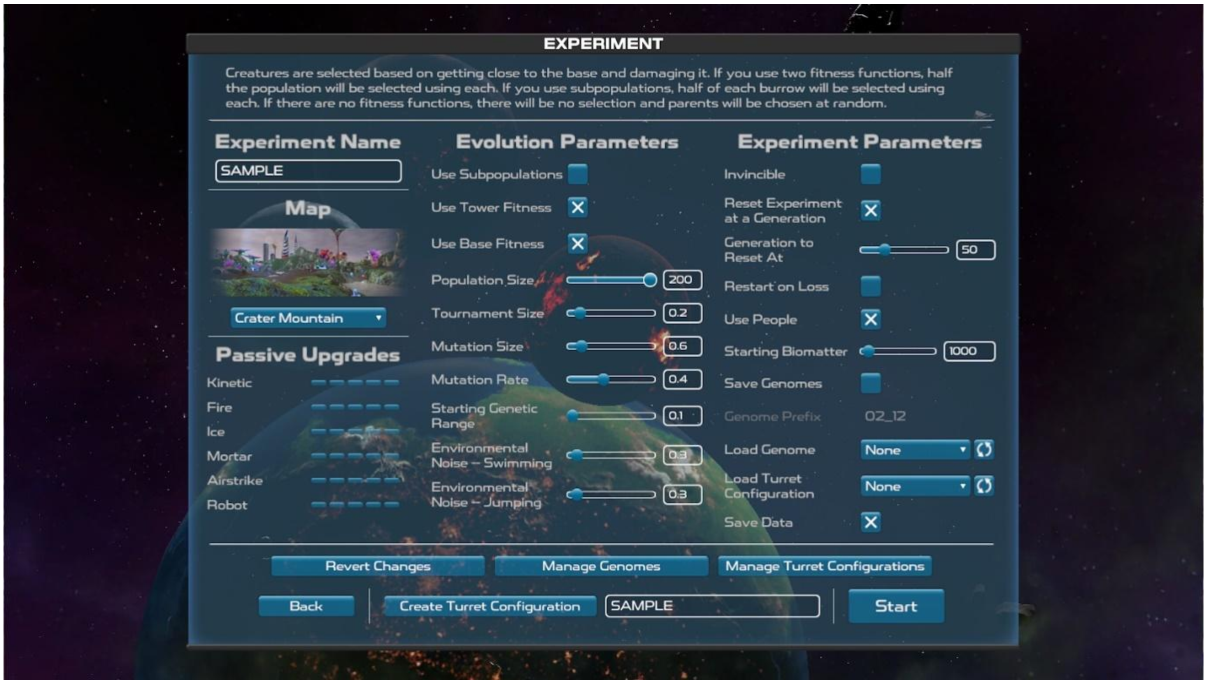


Figure S1. A screenshot of the Experiment Mode user interface.

| Parameter Name | Value Used | Description |
|---|---|---|
| MAP | Crater Mountain | One of 16 game maps |
| PASSIVE UPGRADES | None | Upgrades earned through gameplay |
| USE SUBPOPULATIONS | Off | When ON, causes each Protean nest to act as its own subpopulation. When OFF, all nests belong to one panmictic population |
| USE TOWER FITNESS | Determined by experiment | When ON, Proteans accrue fitness by damaging towers (ON during FitClone and OnlyFit. OFF during OnlyClone and Control) |
| USE BASE FITNESS | Determined by experiment | When ON, Proteans accrue fitness by approaching and damaging the player's base (ON during FitClone and OnlyFit. OFF during OnlyClone and Control) |
| POPULATION SIZE | 200 | The number of Proteans per generation |
| TOURNAMENT SIZE | 0.2 | The proportion of the POPULATION SIZE used in each tournament during the selection phase |
| MUTATION SIZE | 0.6 | The standard deviation of a Gaussian distribution from which mutations are drawn. |
| MUTATION RATE | 0.4 | The per locus probability of an individual Protean acquiring a mutation |
| STARTING GENETIC RANGE | 0.1 | Variation added to genetic values at the beginning of the game |
| ENVRIONMENTAL NOISE - SWIMMING | 0.3 | A value that adds random noise to the genetic value for swimming for each individual. |
| ENVIRONMENTAL NOISE – JUMPING | 0.3 | A value that adds random noise to the genetic value for jumping for each individual. |
| INVINCIBLE | On | When ON, Proteans can damage player structures, but the structures are never destroyed. |

| | | |
|---|---|---|
| RESET EXPERIMENT AT A GENERATION | On | Allows experiment mode to run replicates using the same game settings. |
| GENERATION TO RESET AT | 50 | Allows the user to set the number of generations for each replicate |
| RESTART ON LOSS | Off | Allows the game to restart if the player's base is destroyed |
| USE PEOPLE | Determined by experiment | A Boolean that sets whether civilians are present on the map (ON during FitClone and OnlyClone. OFF during OnlyFit and Control) |
| STARTING BIOMATTER | 1000 | Biomatter is the currency used when playing the game |
| SAVE GENOMES | Off | Allows the game to use previously evolved genomes |
| GENOME PREFIX | NA | Adds a tag to the name of a saved genome |
| LOAD GENOME | NA | Load a previously evolved genome that was saved using SAVE GENOMES |
| LOAD TURRET CONFIGURATION | Determined by experiment | Put a previously created turret arrangement on the map |
| SAVE DATA | On | Save genomic data created during the experiment |
| REVERT CHANGES | NA | Allows the user to reset the Experiment Mode options to their defaults |
| MANAGE GENOMES | NA | Opens the "SavedGenomes" folder in the game's directory on your computer |
| MANAGE TURRET CONFIGURATIONS | NA | Opens the "TurretConfigurations" folder in the game's directory on your computer |
| CREATE TURRET CONFIGURATION | NA | Loads the map without any Proteans to allow the player to freely place turrets |

Table S3. Game parameters that can be adjusted in Project Hastur's experiment mode. The "Value Used" column indicates the associated setting that was used for the experiments in this paper.

For any given replicate, the player was tasked with placing turrets subject to the following constraints: players were required to use exactly 21 autocannon turrets, players could use any number of Energy Matter Converters (EMC) to spread their turrets across the map, and players must place at least one turret by each EMC.

**Text S3. Additional Game Information**

There are four different types of turrets: kinetic (represented by the "Autocannon"), ice (represented by the "Chip shredder"), acid (represented by the missile), and fire (represented by the "Flamethrower"). The four turret types can be seen in Figure S2. For the experiments discussed in this paper, only the Autocannon turret was used. Turrets will only work when placed within a certain radius of an Energy Matter Converter (EMC) or base (Figure S3).

Turrets were placed according to the strategy appropriate for the current replicate. The Burrow strategy focused on protecting areas of the map where Proteans emerge (Figure S4). When employing the Clump strategy, the player created clumps of turrets that the Proteans had to walk past to get to the player's bases (Figure S5). Finally, when employing the People strategy, the player protected areas where civilians walk around (Figure S6).

Civilians are non-player characters (NPCs) who leave buildings to survey the land (Figure S7). They walk along a set path around the buildings, and occasionally crouch down to take samples or simply stand in the same spot for a small amount of time. These civilians are vulnerable to Protean attacks, and clones of the attacking Protean will violently emerge from the attacked civilian. The number of clones that are produced is determined by the size of the attacking Protean.

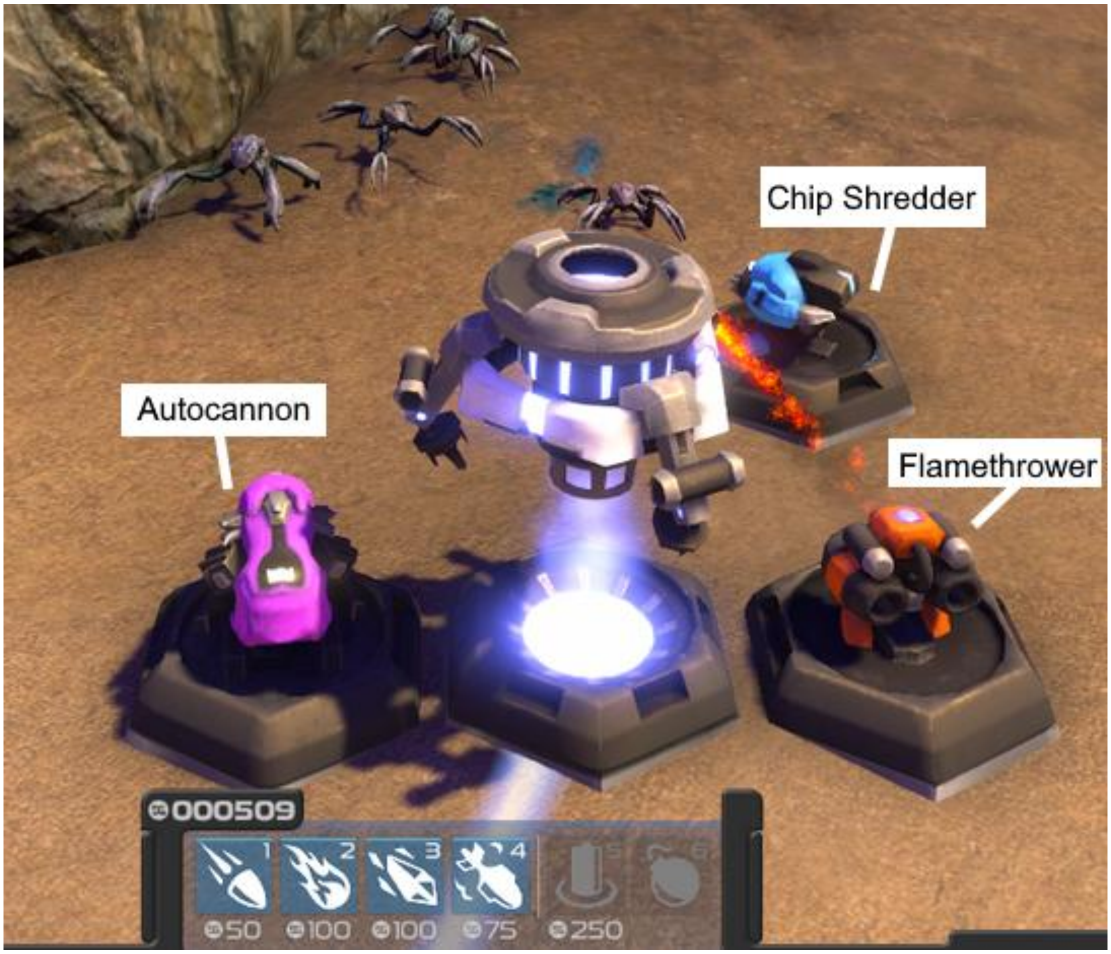


Figure S2. A screenshot of the three different types of turrets: Autocannon, Chip Shredder, and Flamethrower. These three turrets are placed around an Energy Matter Converter (EMC).

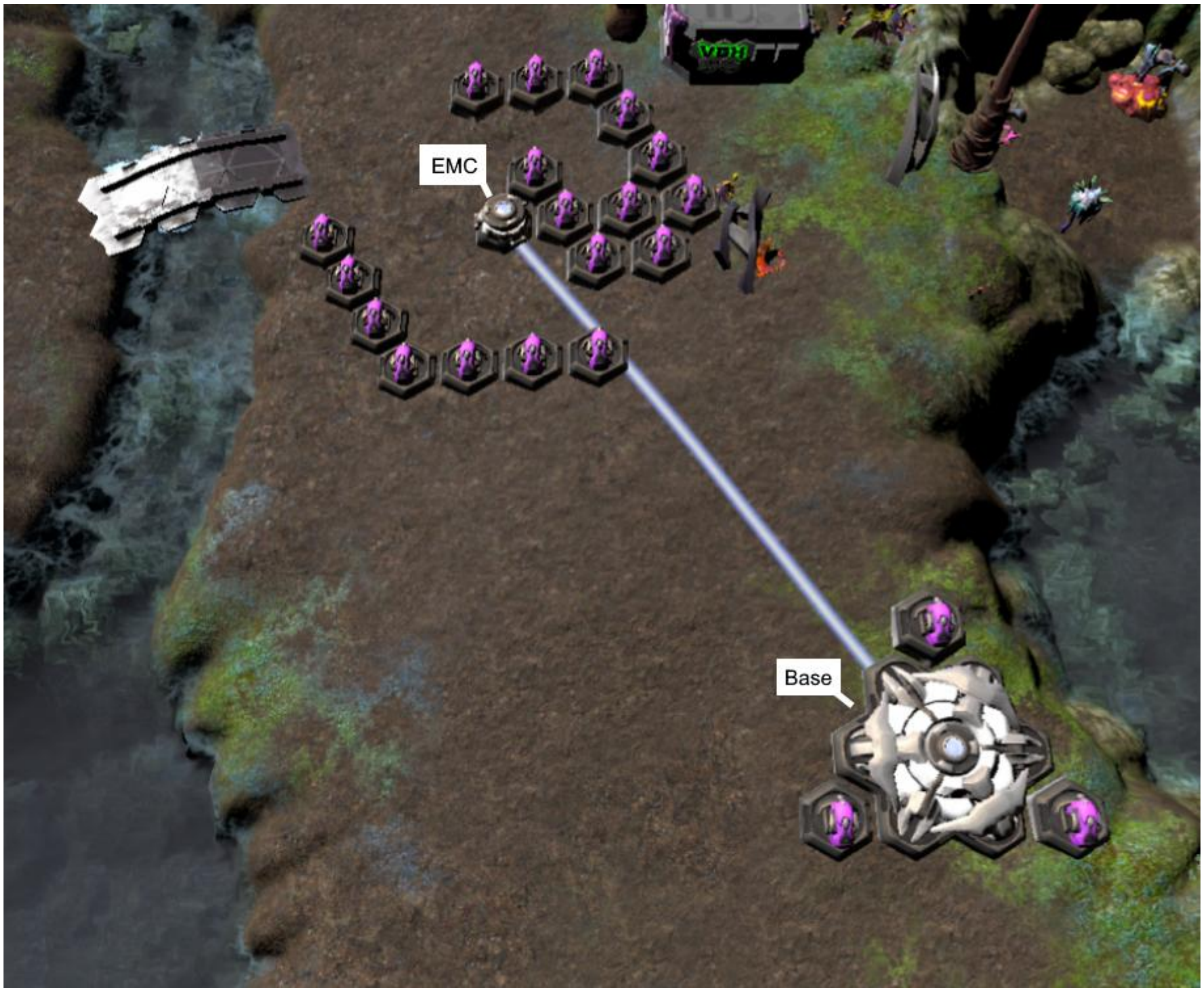


Figure S3. A screenshot depicting an EMC and a Base with several turrets surrounding them.

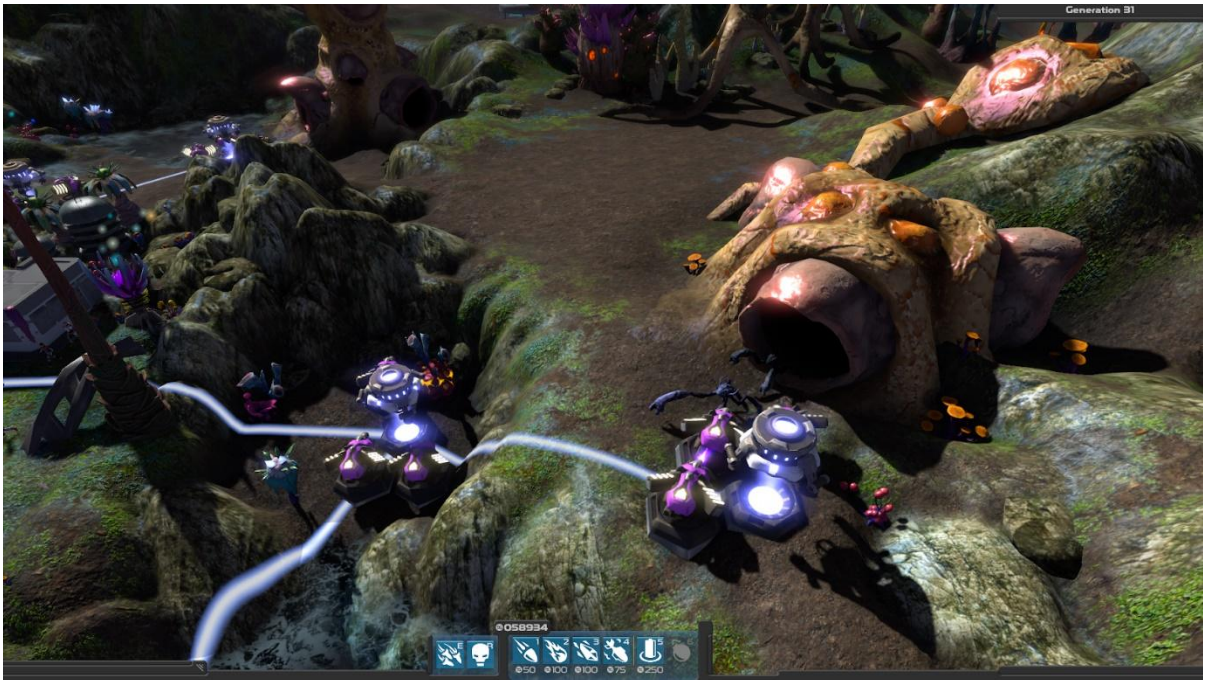


Figure S4. Screenshot of turret positions for the Burrow strategy.

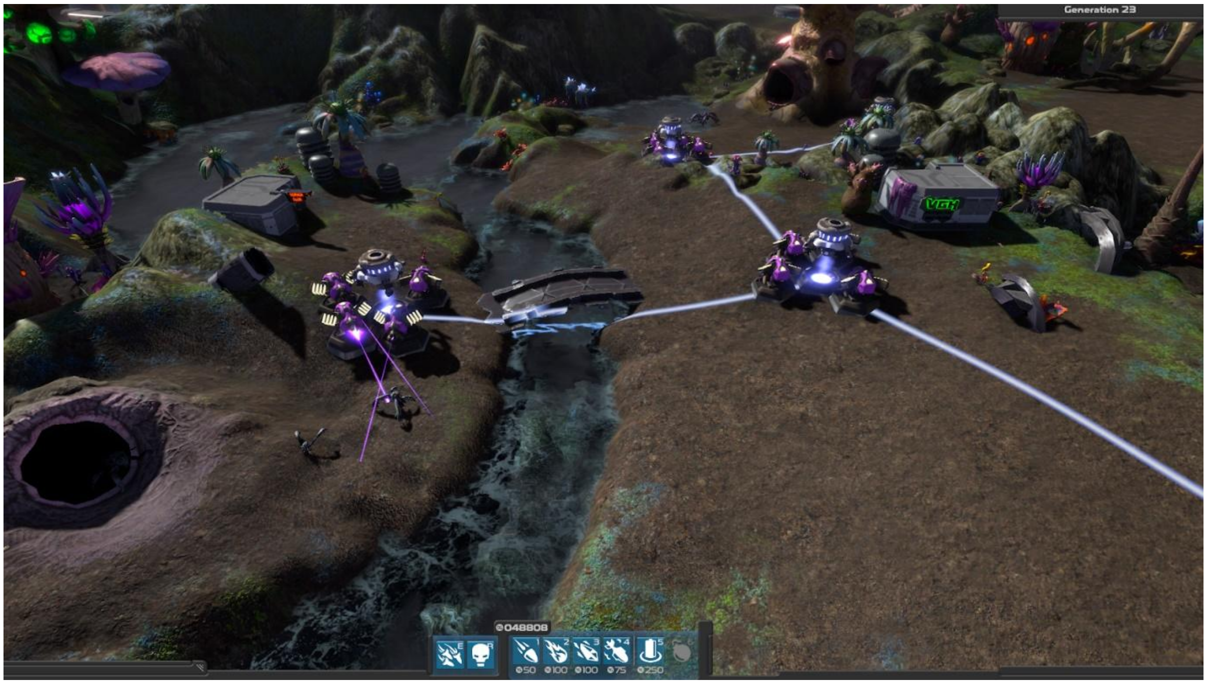


Figure S5. Screenshot of turret positions for the Clump strategy.

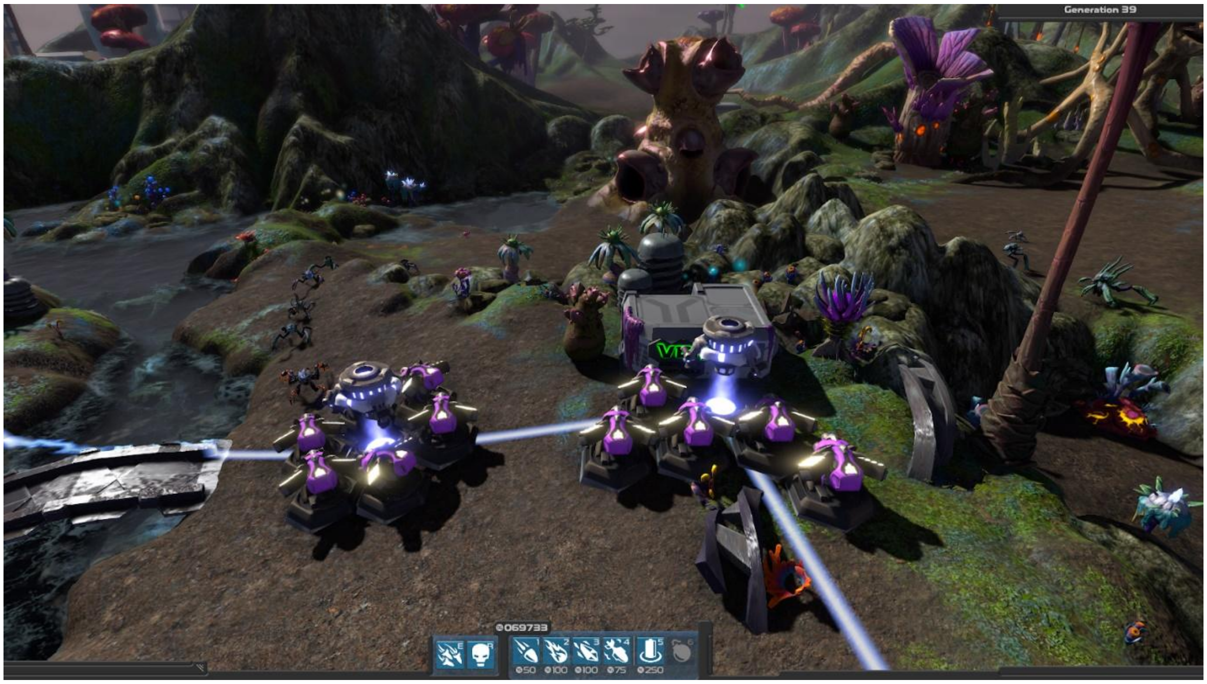


Figure S6. Screenshot of turret positions for the People strategy.

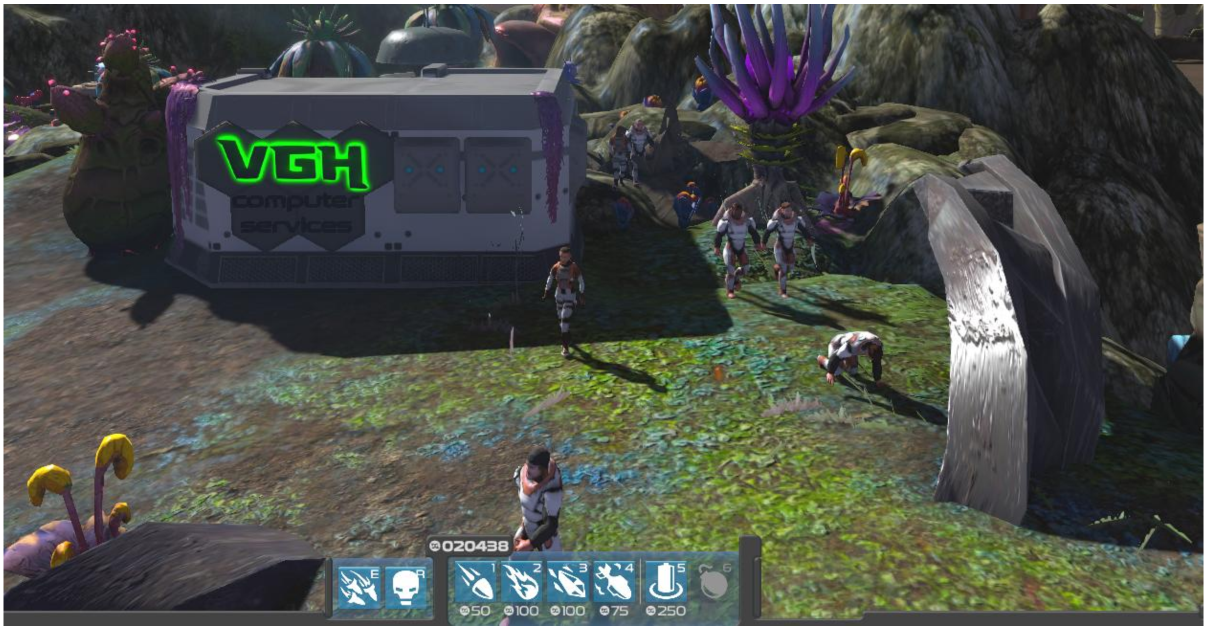


Figure S7. Screenshot of civilian activity around the building from which they emerge.

**Text S4. Dendrogram Creation Details**

In an attempt to visually represent the different kinds of Proteans that evolved in each replicate, we created dendrograms using the centers of clusters created from the final generation's trait data for each replicate. Dendrograms were created by first finding the Euclidean distances between cluster centers and performing hierarchical clustering using the function "hclust()" from the *stats* R package (R Core Team 2021). The resulting hclust object was then used with the function "as.phylo()" from the *ape* R package (Paradis and Schliep 2019) to plot the hclust object as a dendrogram. Figure S8 displays an example dendrogram from the last generation for a random replicate using the Burrow strategy from experimental treatment OnlyClone. The tip labels for this dendrogram are arbitrary names for the clusters based on their order in the distance matrix. We attempted to give these tips more informative names by looking at the trait centroids in each cluster (Table S4). Our initial attempts involved naming the tips based on the highest-ranked traits from each cluster. For example, tip 4 could be named "Fast-Turn Pylon Attacker" (its highest-ranked traits are Walk, Run, Acceleration, TurnRate, and Attraction2), but only naming the tips by the best traits in each cluster ignores the other traits that make Proteans unique. Tip 4's lowest-ranked traits are Health and ColliderSurfaceArea, which in conjunction with the highest-ranked traits, leads to the conclusion that the Proteans in that cluster preferentially evolve ways to escape turret fire instead of evolving ways to deal with getting hit. With both the excellent and terrible traits in mind, this Protean could be named "Glass Fast-Turn Pylon Attacker" to account for its low health. This method of naming, while tedious, makes logical sense when considering just one replicate at a time. However, there are different ways to be a "Glass Fast-Turn Pylon Attacker" that emerge in other replicates. A Protean with that same name could have highly-ranked Walk, Run, TurnRate, and Attraction2 traits, but maybe Acceleration is not its strong suit. Perhaps we could incorporate the lack of Acceleration into a new name like "Glass Moderately-Fast-Turn Pylon Attacker." Ultimately, the farther we go into creating specific names to describe each cluster, the more uninformative the task becomes because the vast majority of clusters across our entire dataset would have unique names.

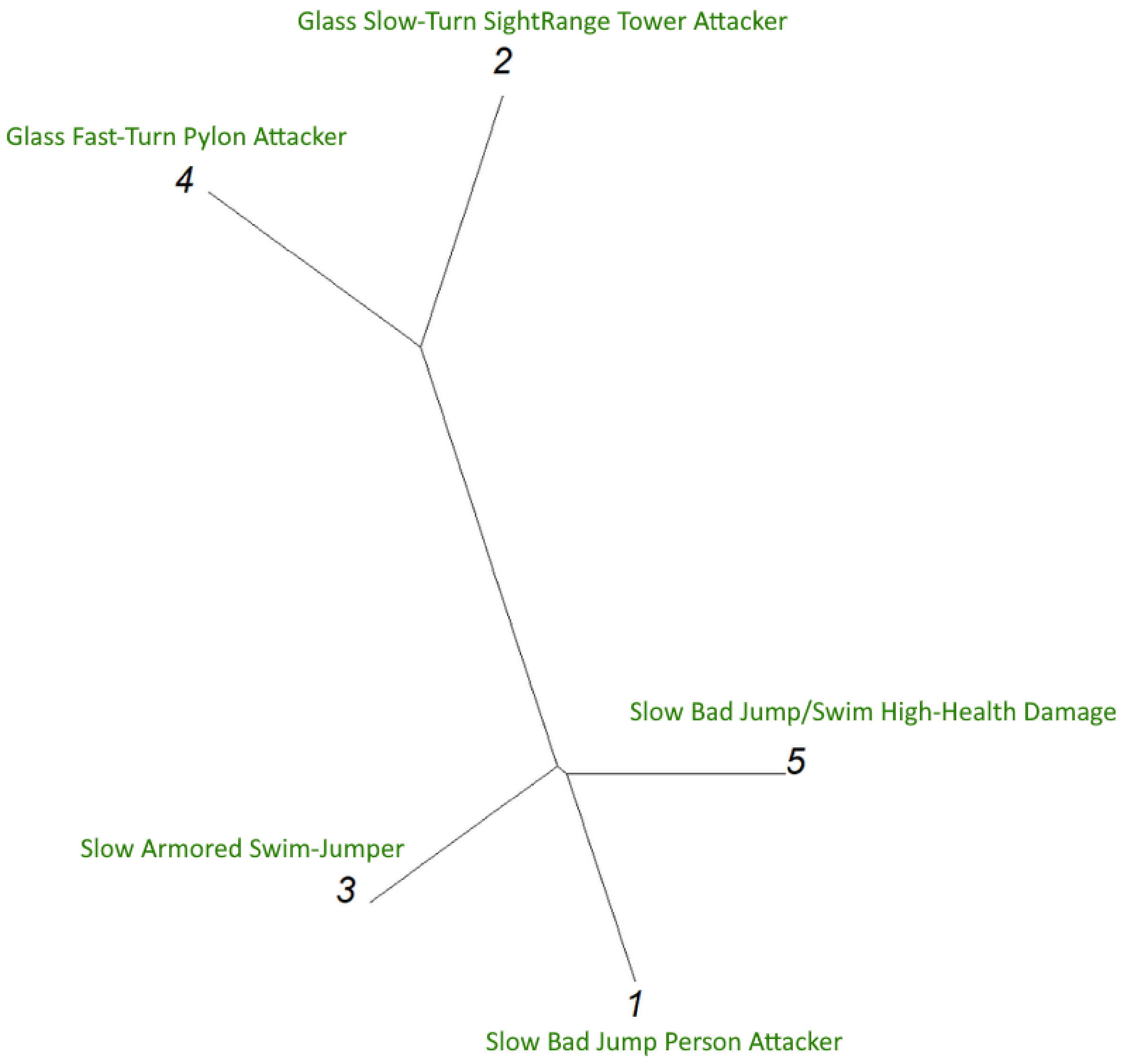


Figure S8. Example dendrogram for the last generation for a replicate from experimental treatment OnlyClone using the Burrow strategy. Potential names for each tip are written in green.

| Trait | Description 1 | Description 2 | Description 3 | Description 4 | Description 5 |
|---|---|---|---|---|---|
| Health | Average health | BAD rank health | GOOD rank health | BAD rank health | GREAT rank health |
| SightRange | Average sight | GREAT rank sight | BAD rank sight | GOOD rank sight | BAD rank sight |
| Armor | Average armor | BAD rank armor | GREAT rank armor | GOOD rank armor | BAD rank armor |
| Damage | GOOD rank damage | BAD rank damage | BAD rank damage | Average damage | GREAT rank damage |
| WalkSpeed | Average walk | GOOD rank walk | BAD rank walk | GREAT rank walk | BAD rank walk |
| RunSpeed | Average run | BAD rank run | GOOD rank run | GREAT rank run | BAD rank run |
| Acceleration | Average accel | GOOD rank accel | BAD rank accel | GREAT rank accel | BAD rank accel |
| TurnRate | GOOD rank turn | BAD rank turn | BAD rank turn | GREAT rank turn | Average turn |
| Swim | Average swim | BAD rank swim | GREAT rank swim | GOOD rank swim | BAD rank swim |
| Jump | BAD rank jump | GOOD rank jump | GREAT rank jump | Average jump | BAD rank jump |
| ColliderSurfaceArea | GREAT rank collider | BAD rank collider | GOOD rank collider | BAD rank collider | Average collider |
| Attraction0 | BAD rank at0 | GREAT rank at0 | Average at0 | GOOD rank at0 | BAD rank at0 |
| Attraction1 | GREAT rank at1 | BAD rank at1 | BAD rank at1 | GOOD rank at1 | Average at1 |
| Attraction2 | Average at2 | BAD rank at2 | BAD rank at2 | GREAT rank at2 | GOOD rank at2 |

Table S4. Example cluster description matrix created from ranking traits between clusters. Each column represents a cluster, each row represents a trait, and each cell describes the ranking of that trait compared to the other clusters.
